# Detecting co-selection through excess linkage disequilibrium in bacterial genomes

**DOI:** 10.1101/2023.08.04.551407

**Authors:** Sudaraka Mallawaarachchi, Gerry Tonkin-Hill, Anna K. Pöntinen, Jessica K. Calland, Rebecca A. Gladstone, Sergio Arredondo-Alonso, Neil MacAlasdair, Harry A. Thorpe, Janetta Top, Samuel K. Sheppard, David Balding, Nicholas J. Croucher, Jukka Corander

## Abstract

Population genomics has revolutionised our ability to study bacterial evolution by enabling data-driven discovery of the genetic architecture of trait variation. Genome-wide association studies (GWAS) have more recently become accompanied by genome-wide epistasis and co-selection (GWES) analysis, which offers a phenotype-free approach to generating hypotheses about selective processes that simultaneously impact multiple loci across the genome. However, existing GWES methods only consider associations between distant pairs of loci within the genome due to the strong impact of linkage-disequilibrium (LD) over short distances. Based on the general functional organisation of genomes it is nevertheless expected that the majority of co-selection and epistasis will act within relatively short genomic proximity, on co-variation occurring within genes and their promoter regions, and within operons. Here we introduce LDWeaver, which enables an exhaustive GWES across both short- and long-range LD, to disentangle likely neutral co-variation from selection. We demonstrate the ability of LDWeaver to efficiently generate hypotheses about co-selection using large genomic surveys of multiple major human bacterial pathogen species and validate several findings using functional annotation and phenotypic measurements. Our approach will facilitate the study of bacterial evolution in the light of rapidly expanding population genomic data.

## Main

The rapid rate of evolution of bacterial genomes has made them a popular target of studying selection in both experimental and natural populations. The emergence of affordable highresolution population genomics a decade ago ushered us into a new era with improved possibilities to investigate both microscopic and macroscopic evolution of bacterial genomes, including for example codon bias [1], intergenic selection [2], and variation in gene content [3]. Despite steady progress in genome-wide association study (GWAS) methodologies specifically designed for bacterial populations [4,5], the difficulty of measuring quantitative phenotypic variation in large numbers of isolates has restricted the use of GWAS mostly to the study of antibiotic resistance, with a few exceptions covering for example duration of colonisation [6], genome-wide transcriptomics [7] and virulence [8]. Traits such as reproductive rate, survival and transmissibility are of key interest in bacteria but remain difficult to measure in sufficient numbers in natural populations. Motivated by this obstacle, phenotype-free approaches to uncovering signals of selection have been introduced [9–12], based on the rationale that positively selected variation in complex traits would likely be caused by synchronised changes in multiple genes and/or regulatory elements that are detectable from excess linkage disequilibrium (LD) between distant sites across a sample of genomes. This provides a methodological toolkit complementary to GWAS enabling analyses termed genome-wide epistasis and co-selection studies (GWES), which has been recently used to unravel signals of selection due to epistasis for a wide diversity of bacteria [13–16]. Arnold et al. demonstrated that under selection, even relatively weak epistasis is sufficient for driving adaptation in fairly recombinogenic bacteria, which provides a more theoretical justification for the GWES principle [17].

A major limitation of the existing GWES approaches is that their scope is limited to discovering links between distant loci within the region where LD asymptotes towards its lower bound. However, many co-evolving loci are organised into clusters of genes (e.g. cotranscribed in operons). Therefore, studying only long-range links ignores most of the allelic co-variation occurring in genomes. A fine-scale haplotype structure analysis of *Neisseria gonorrhoeae* indeed revealed numerous likely examples of positive co-selection in different regions of the chromosome of this highly recombinant species, and extensive population genetic simulations suggested that such LD patterns could be explained by either directional selection on horizontally acquired alleles, or balancing selection maintaining the diversity [18]. Motivated by these insights, we aimed at developing a scalable statistical approach that enables disentangling neutral LD from co-selection and epistasis at any genomic distance.

GWES methods generally exploit the decay of LD as a function of genomic distance to label SNP pairs as ‘outliers’ with respect to the background distribution of LD estimated from population data. Similar to Arnold et al. [18], it is possible to extend the notion of ‘outlier’ level LD to SNPs in close proximity to each other by simulating the distribution of LD strength as a function of base pair distance using a neutral Wright-Fisher model with suitable parameters governing mutation and recombination events. These simulations approximate the population level co-variation of alleles expected under neutrality and can be used to screen pairs of loci for outliers indicating co-selection. We show that this approach works well and maintains a low false positive rate. However, as fitting of the neutral model parameters and forward simulation of the fitted model in a sufficiently large number of replicates is computationally costly, we developed an empirical model-free approximation that is scalable to large population genomic datasets. The model-free method is motivated by the common assumption that a majority of the observed LD within a bacterial population reflects near-neutrality [19–22], and therefore it is feasible to use the empirical LD decay distribution for calling outliers. Moreover, the approach accounts for heterogeneity in evolutionary rates, such as mutation and recombination hotspots.

The model-free algorithm is implemented as an open-source R package ‘LDweaver’ (https://github.com/Sudaraka88/LDWeaver), which can be used to perform a comprehensive GWES in large-scale bacterial datasets. LDWeaver incorporates the functionality of the popular GWES package Spydrpick for long-range LD outlier detection [12] and extends this by allowing analysis of LD at any genomic distance. LDWeaver provides automated functional annotations on all putative co-selected SNPs and generates an array of visualisations to allow users to efficiently explore the results. We use published population genomic data for the major human pathogens *Streptococcus pneumoniae* [23,24], *Campylobacter jejuni [25], Escherichia coli [26]* and *Enterococcus faecalis [27]* to identify both known and novel signals of co-selection linked to the molecular basis of pathogenicity, survival and other key bacterial phenotypes.

## Results

### Overview of LDWeaver

Performing GWES analysis using LDWeaver requires two inputs, a multiple sequence alignment (MSA) and the annotation file of the reference in Genbank or Gff3 format. Following SpydrPick [12], LD between each SNP pair is measured using mutual information (MI). Generally, short-range links (i.e., SNPs in genomic proximity) are in much stronger LD compared to the long-range (see Figure 1a). Therefore, to determine a threshold for outlier calling, it is necessary to model the decay in LD with genomic distance. The shape of this decay is determined by factors such as the type of species, population structure, local mutation and recombination rates, variation in gene content and selection pressures. To account for this heterogeneity, LDWeaver measures the per-site mean Hamming distance within coding regions around the chromosome and clusters them based on this estimated local diversity (see Figure 1b). The background decay, modelled separately for each cluster, is used to estimate the 95th percentile of MI values at each region or base pair-separation (bp-sep) (see Figure 1c-e). Since long-range background LD is uniform (see Figure 1f), the Tukey outlier approach in SpydrPick is sufficient. After determining the local thresholds and calling outliers, LDWeaver provides a ranked list of all loci pairs that are under putative coselection.

**Figure 1.**
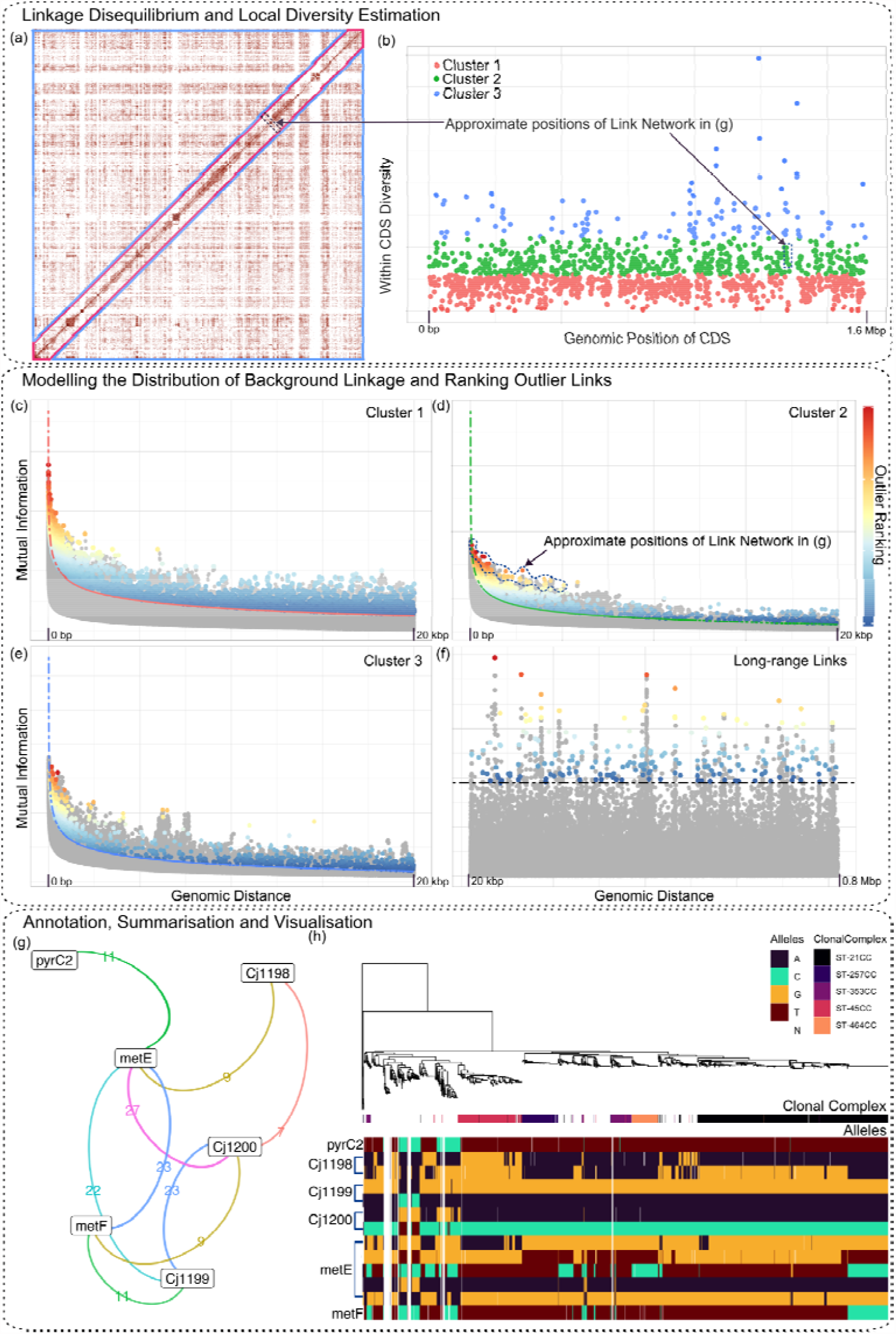
Overview of the LDWeaver pipeline. (a) Genome-wide linkage disequilibrium (LD) for 1,480 *Campylobacter jejuni* isolates [25]. LD is measured using mutual information (MI) and a weighting strategy is employed to account for population structure. The axes correspond to genomic positions in the NCTC 11168 reference genome and the colour intensity reflects the strength of LD. Blue triangles outline regions of long-range LD, while the short-range high-LD region is outlined in red. (b) Genomic diversity within each coding region (CDS) is used to account for local variation in short-range background LD. Each point corresponds to a CDS and the vertical axis shows the average number of sites that differ from the reference. K-means clustering is used to divide the CDS into 3 clusters (see legend). Sites from intergenic regions are allocated to the nearest cluster. (c), (d), (e) and (f) GWES Manhattan plots show the modelled background LD (dashed coloured lines) and outlier ranking of SNP pairs based on significance level (colour bar in the rightmost panel) for short-range links in the three clusters (c, d and e) and for long-range links (f). The plot shows the ranked MI values for SNP pairs against genomic distance. Links that are either estimated as indirect or with MI below the background LD are depicted in grey and not ranked. In (f) the background LD is invariant to genomic distance and computed using the Tukey criteria (dashed black line). (g) The LDWeaver network plot generated for *metE* summarises all the outlier links involving a site in the gene. Here, the edges are coloured based on the number of links between linked genomic region nodes (see *Campylobacter* results section). (h) Investigating the allele distribution within the linked region (Alleles panel) shows that several deletions and mutations observed within several clonal complexes are driving the high LD signal picked up by LDWeaver.

LDWeaver includes several options to ease the task of generating a list of potential epistatic SNP-pairs for expert manual curation and wet lab validation. First, all outlier links are annotated using SnpEff [28] and links that include a non-synonymous mutation are given a higher priority. LDWeaver summarises links into networks (see Figure 1g), which helps to prioritise the most promising genome regions. Finally, LDWeaver can be used to visualise the allele distributions within these networks (see Figure 1h).

### LDWeaver detects co-evolutionary links in multiple pathways in *S. pneumoniae*

The analysis of pneumococci focussed on two populations isolated from carriage in contrasting settings: Mae La, Thailand and Massachusetts, USA. The Mae La sample comprised 2,663 high quality assemblies (accessions available in supplementary information) [29] collected from mother and infant pairs [30]. The Massachusetts sample [24] comprised 616 draft genomes (accessions available in supplementary information) of similar quality [29]. Both datasets were aligned to the ATCC 700669 (accession code FM211187) reference genome [31], and after filtering out sites with minor allele frequency (MAF) < 0.01 and gap frequency > 0.15, respectively 88,603 and 89,386 SNPs were retained for analysis (see supplementary Figs. 1 and 2 for LDWeaver plot panels).

Long-range interactions included strong signals of co-evolution between *pbp2b* and *pbp2x* in both populations. These genes are key in determining resistance to beta lactam antibiotics and these interactions were reported previously [12]. Another interaction conserved across both populations was that between three loci encoding immunogenic surface-exposed degradative enzymes [32]: the beta-galactosidase BgaA, the immunoglobulin A protease ZmpA, and PabB, encoded by a gene directly downstream of that for the ZmpA paralogue, ZmpB. These co-evolutionary signals may arise through direct interactions on the surface, or indirect effects emerging through immune selection for particular combinations of antigens [33].

The shorter-range interactions were more reproducible across the populations. Adjacent genes functioning in the same metabolic pathway included links consistently identified between the neighbouring coding sequences cdsH (SPN23F08030) and thiI (SPN23F08040), which are both involved in thiamine biosynthesis [34]. Similarly, the adjacent genes SPN23F02450 and SPN23F02460 both encoded proteins predicted to function as N-acetyltransferases. Signals were also identified in both populations between the overlapping genes SPN23F08250 and SPN23F08260, encoding subunits of a transporter of unknown function [35]. Hence, LDweaver can identify signals of proteins likely to be co-evolving as participants in the same metabolic pathways.

Another regulatory locus involved in short range interactions across both datasets is that encoding the competence regulator TfoX. Multiple sites were in strong LD in the subset of isolates encoding this locus, which was absent in a minority of isolates (see Fig. 2a). Yet these sites are not in the gene itself, but in the flanking intergenic regions, or the adjacent *alaDH* pseudogene that encodes an apparently functionally unrelated alanine dehydrogenase. This suggests that these paired sites do not interact functionally. A search for the functional sequence of the *alaDH* gene identified intact versions in other streptococci. Further alignments indicated that the *tfoX*-*alaDH* pairing was intact in *Streptococcus anginosus*. Hence, the variable distribution of this locus in *S. pneumoniae* is likely the consequence of one, or more, interspecies transfers through homologous recombination introducing the gene cassette into pneumococci, followed by degradation of the *alaDH* gene into a non-functional form [36]. Hence the elevated LD identified in this case may be the consequence of a relatively recent introgression [37] from another streptococcal species, demonstrating LDWeaver can identify loci under sufficiently strong selection to drive interspecies transfers [36].

**Figure 2.**
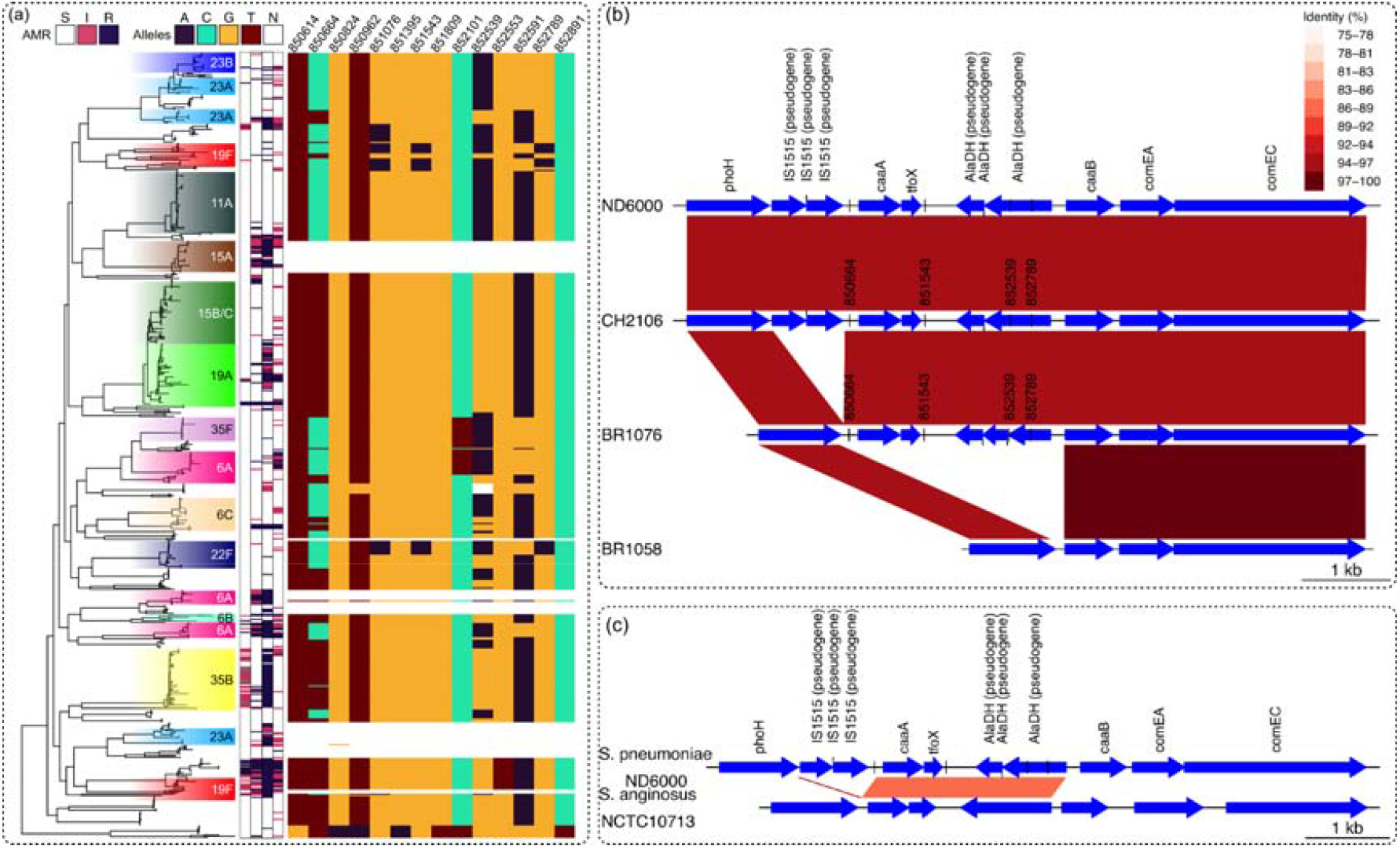
Overview of genomic variations of the flanking region of the TfoX competence regulator demonstrated using the Massachusetts S. pneumoniae dataset. (a) The phylogenetic tree (n = 616) is coloured according to the serotype shown to the right of the tree. The 4 bars immediately to the right denote the antimicrobial resistance data for ceftriaxone, erythromycin, benzyl penicillin and trimethoprim, respectively. The key above indicates the colour shadings for S - sensitive, I - intermediate and R - resistant. Allelic variation in the tfoX locus is shown by the rightmost heatmap and the key above shows the colour for each nucleotide with N indicating an ambiguous base. SNP positions above the heatmap are based on the ATCC 700669 reference. This suggests that the tfoX locus is present in most pneumococci, and can be divided into three genotypes: that containing the major alleles at each polymorphic site; that containing the minor alleles at sites 850664 and 852539, and that containing the minor alleles at these two sites, as well as sites 851543 and 852789. (b) The alignment of representatives of the four observed genotypes in the dataset, demonstrated here using sample data from selected isolates. Colour shading indicates the identity between regions. In ND6000 (ERR129187) and CH2106 (ERR129095), the flanking region is intact with multiple insertion sequences, two functional genes, a non-coding region and an alaDH pseudogene. The locus is intact, but the insertion sequences are not observed in BR1076 (ERR129048). In contrast, the entire locus is missing in BR1058 (ERR129043). (c) Shows the alignment between FM211187 and S. anginosus NCTC10713 genomes. For the region of interest, this is the most similar locus among streptococcal species. Despite the general divergence between S. pneumoniae and S. anginosus, the tfoX-alaDH gene pair is intact in both species with high similarity (same key as in (b) to show region identity). However, the dissimilarity between the flanking regions demonstrates the typical level of divergence between the genomes. Hence, the localised similarity indicates a possible recent introgression into the pneumococcus.

Another transporter (encoded by SPN23F03500) was linked to an adjacent pseudogene (SPN23F03510) in both populations (see supplementary Fig. 3). The undisrupted sequence of SPN23F03510 is predicted to function as a lantibiotic synthesis protein, suggesting the associated transporter is likely to be a self-immunity protein. Hence this pairing likely represents an example of a non-producing, immune-only “cheater” bacteriocin phenotype [38], common in pneumococci [39]. The most variable bacteriocin-encoding locus in the pneumococcal genome is the *blp* gene cluster [40], in which multiple interactions between genes were detected. These included interactions between the genes encoding the BlpRH quorum sensing two component system, and the gene encoding the cognate peptide pheromone, BlpC [41], identified in both populations. Further links were found in the Massachusetts sample only, which likely represent relationships between the synthesised bacteriocins and corresponding immunity proteins [40].

### LDWeaver Identifies patterns of co-evolution of cytolethal distending protein toxin subunits in *C. jejuni*

*C. jejuni* is a leading cause of food-borne bacterial gastroenteritis worldwide, associated with the consumption of contaminated poultry meat. It is well-adapted to colonise the gut of the majority of mammalian and avian host species and has been isolated from many environmental sources. The successful colonisation of strains is often host-specific for the majority of lineages (specialist clonal complexes) and this is reflected in the population structure during phylogenetic comparison of genomes. Certain clonal complexes (CCs) are also known to be host generalists, where the same lineage is well-adapted to colonise and to survive in multiple different hosts.

The population of *C. jejuni* is structured by well-defined, genetically similar clusters of isolates (clonal complexes) which are documented to be maintained over time [25], despite the very frequent homologous recombination occurring across the known lineages [25,42]. This high frequency of recombination, which is not limited to any particular region of the genome, would generally break down the LD observed in *C. jejuni* lineages and therefore, potentially disrupt co-selected/epistatic functional groups of genes throughout the genome. Despite the high recombination, the population structure has remained stable over long periods of time making *C. jejuni* an excellent candidate species for the analysis of short- and long-range epistasis and co-selection links.

A collection of 1,480 previously published *C. jejuni* genomes [25] consisting of 18 different human, animal and environmental sources and 37 CCs were selected for LDWeaver analysis (accessions available in supplementary information). These were aligned against the NCTC 11168 reference genome [43] using snippy. After removing sites with MAF < 0.01 and gap frequency > 0.15, 102,591 SNPs were retained for LDWeaver analysis (see supplementary Fig. 4 for the LDWeaver plot panel).

Analysis of the 250 most statistically significant short-range interactions identified strong signals of co-evolution between genes with functions mostly related to virulence such as: amino acid ABC transporters, flagellar biosynthesis, periplasmic and outer membrane proteins, antibiotic efflux genes, among others. One of the most promising findings was multiple highly-significant links located within the cytolethal distending toxin (CDT) genomic region. The CDT is a protein toxin composed of three subunits: CdtA, CdtB and CdtC encoded by a *cdtABC* operon, and is considered one of the most important virulence factors for *Campylobacter* pathogenesis. CDT acts to halt host cell division by cell cycle arrest at the G_2_ stage occurring before mitosis [44]. CdtA and CdtC are anchored into the membrane and act to deliver CdtB to the host cell which arrests the cell cycle. CdtB has the toxin activity but is reliant on CdtA and CdtC for its binding and delivery to the host cell [45].

Our results showed that the three subunits were well conserved throughout the dataset, except for three distinct clusters of isolates identified in wild birds (see Figure 3). There was high variability of the presence/absence and allelic variation of the three CDT subunits across these three clusters of isolates. For example, all three subunits were absent from the wild bird-associated ST-1287 CC. Another wild bird-associated clade, ST-1034 CC (mixed) consisted of synonymous/non-synonymous nucleotide changes in all subunits compared with other sources. Finally, isolates belonging to the third wild bird-associated clade, ST-1325 CC consisted of a mix of both the same synonymous/non-synonymous nucleotide changes as seen in ST-1034 CC, some of the isolates in this cluster also exhibited the same variation as observed in the other sources and CCs within the dataset, while some isolates had an absence of various CDT subunits (see Figure 3). A recent study comparing the gene sequences of the *cdtABC* operon of wild bird, broiler chicken and human sources [46] confirms the LDWeaver-generated hypothesis of significant co-selection occurring within this operon. The study identified high variability of *cdtABC* alleles in wild birds with several alleles producing no functional CDT. Sequence conservation outside of wild bird sources, such as broiler chickens and humans, was also observed, suggesting that the variation of the *cdtABC* operon may play a role in the host range of *Campylobacter* [46].

**Figure 3.**
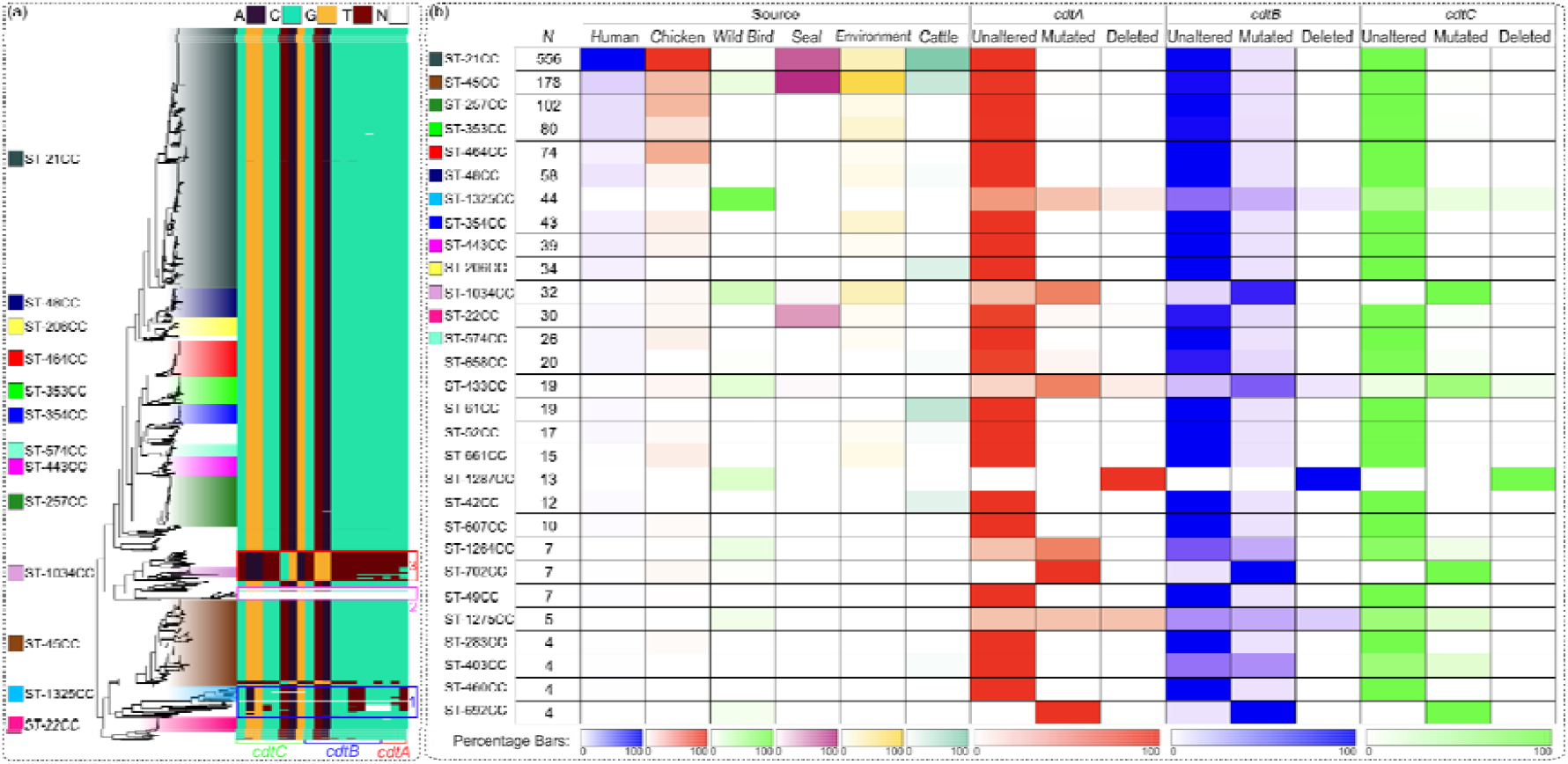
Overview of allelic variation of CDT subunits in the *C. jejuni* dataset. (a) The phylogenetic tree (n = 1,480) is coloured according to the clonal complex and the key to the left of the tree is labelled according to the corresponding topology of each clonal complex. Allelic variation of the three CDT subunits (CdtA, CdtB and CdtC (*x axis*)) encoded by the *cdtABC* operon across the tree is represented by the heatmap to the right of the tree. (b) The association of a particular clonal complex with a source is highlighted by the first panel. The higher the shading (explained by percentage keys at the bottom of the panel), the higher the percentage of isolates belonging to a particular source. Clonal complexes (rows) are ordered by abundance in the *C. jejuni* dataset. Three panels to the right represent the allelic variation within CdtA (red), CdtB (blue) and CdtC (green) subunits. The shading represents the percentage of isolates associated with a particular clonal complex/source with either: 1) the unaltered allele; 2) a nucleotide change (mutated); or, 2) a deletion of the CDT subunit. Shaded keys at the bottom of the three panels represent the percentage of isolates.

Potential co-selection was also identified in the gene cluster containing genes *metE* and *metF* (see Figure 1). These genes are located on an operon involved in methionine synthesis which has been proven to have a vital role in the colonisation of *C. jejuni* to the gastrointestinal tract of different hosts (Kelley et al., 2021). Similar patterns of the presence/absence and mutated versions of this locus were observed within the same wild bird clusters as the *cdtABC* operon (Figure 1h, Figure 3) while also remaining relatively conserved in the remaining sources and CCs.

### LDWeaver recapitulates co-evolving links involved in clade evolution and success of the *E. coli* pandemic clone ST131

To investigate epistasis and co-selection in *E. coli*, we considered a dataset consisting of 2,156 ST131 genome assemblies (accessions available in supplementary information) aligned against the EC958 reference genome [47] using Snippy. Loci with MAF >= 0.01 and gap frequency < 0.15 were included, leading to 44,092 SNPs (see supplementary Fig. 5 for the LDWeaver plot panel). It is noted that the E. coli dataset had the highest amount of LD among all analysed datasets, and the asymptote of supplementary Fig. 5(b) cluster 1 has the largest intercept among all LDWeaver panel plots.

The *E. coli* ST131 lineage belongs to the phylogroup B2 that emerged globally around 20 years ago and is associated with urinary tract (UTI) and bloodstream infections (BSI) [48– 50]. Large epidemiological studies have identified major differences in the virulence and antibiotic resistance between the three main clades of ST131 (A, B and C) [51]. Clade C, which is associated with fluoroquinolone resistance arising from mutations in *gyrA* and *parC* has further been split into two sublineages (C1 and C2) with a distinct pattern of mobile genetic elements (MGEs) and associated antimicrobial resistance (AMR) genes [52,53].

The top-ranking long-range loci pairs in *E. coli* ST131 corresponded to clade C specific SNPs differentiating sublineages C1 and C2 [54]. These included links between sites in *sbmA* (EC958_0513), a transporter involved in the internalisation of peptide antibiotics into the cytoplasm [55] and identified as a virulence factor in avian extra-intestinal *E. coli* (APEC) [56], and sites in i) *nikA* (EC958_3870), a periplasmatic protein from the ATP-binding cassette type nickel transport system acting as the initial receptor of nickel [57], ii) *acrF* (EC958_4822) encoding an efflux pump with homology to the major pump AcrB [58], iii) *lepA* (EC958_2875) encoding a conserved GTPase with a role in the initiation phase of translation [59], and iv) *iscS* (EC958_2841), encoding a cysteine desulfurase implicated in the activity of a number of Fe-S proteins [60]. These clade-specific SNP pairs are spread across the *E. coli* chromosome, separated by at least 1 Mbp, and may have contributed to the recent expansion of the *E. coli* sublineage C2.

The genome wide distribution of LD estimated from LDWeaver revealed multiple interesting patterns (see Fig. 4). For example, the short-range co-evolving SNP pairs highly ranked by LDWeaver correspond to regions involved in the synthesis of the *E. coli* capsule polysaccharide (*kpsM* EC958_3343, *kpsC* EC958_3337, *kpsS* EC958_3338) and type II secretion system located downstream in the chromosome (*gspL* EC958_3345, *gspM* EC958_3344). The observed tight linkage in the *E. coli* capsular region might be critical for having a functional system since the capsule plays an important role as a major virulence factor contributing to the colonisation of different eukaryotic host niches, reducing the efficacy of the immune system by complement inactivation and shaping the horizontal gene transfer mediated by MGEs [61–63]. These results indicate that the variation within the capsule region is also linked to SNPs present in the conserved type II secretion system. Variation in these regions could thus alter the capsule expression in *E. coli*, contributing to a non-capsulated state that allows the introduction of a new pool of MGEs [63].

**Figure 4.**
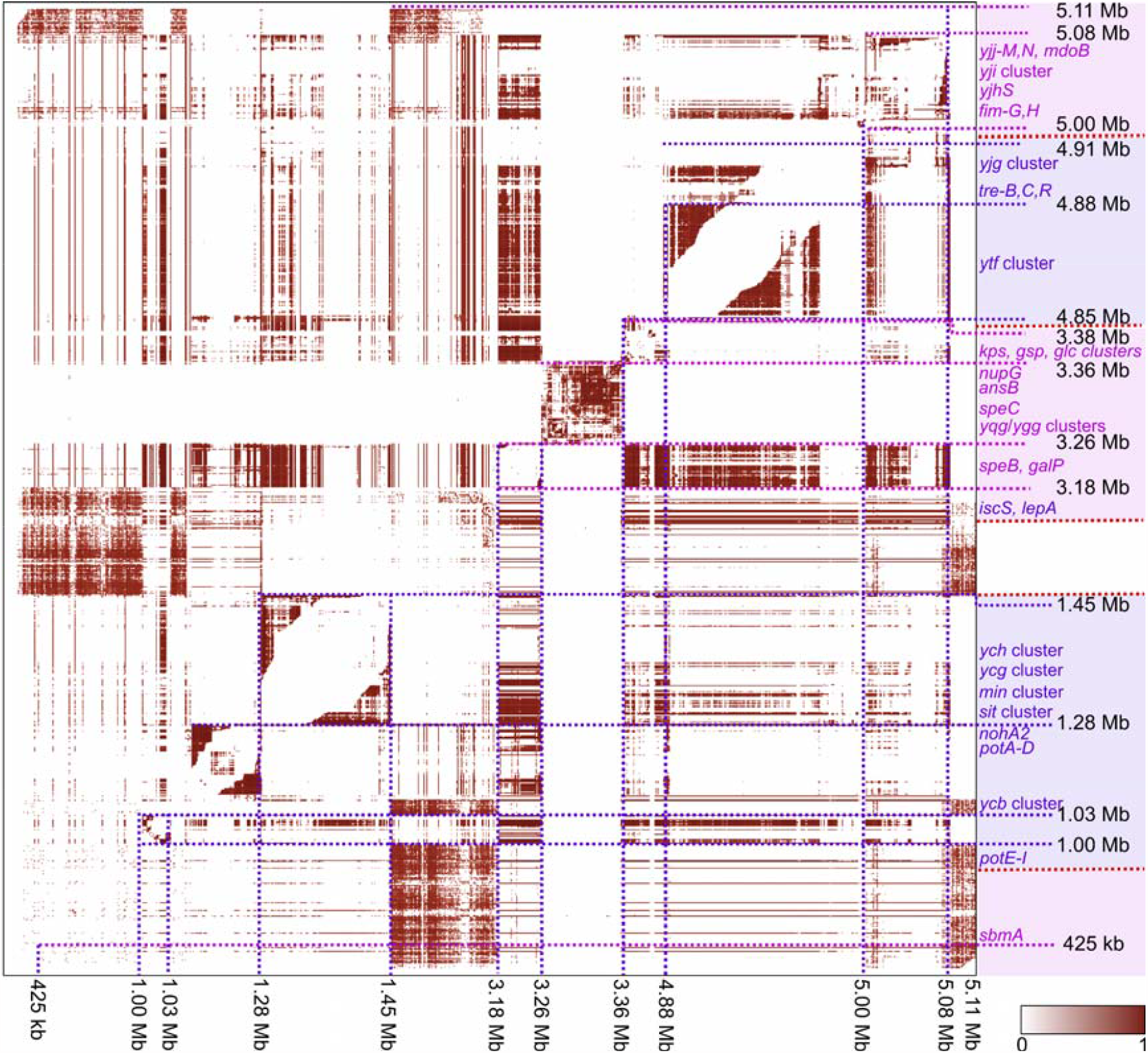
Genome-wide distribution of excess LD in the *E. coli* ST131 dataset (n = 2,156) measured using LDWeaver. Approximate genomic positions are marked to the right and bottom of the LD map as per the EC958 reference genome. Brown shading indicates the amount of LD between sites (see key at bottom right for MI values scaled between 0-1). Entire genomic regions without excess LD are dropped from the figure to enhance the visibility of variation within high LD regions and each new region is coloured in an alternating shade of blue and purple in the right panel. Additionally, the right panel shows genomic regions comprising short-range excess-LD links.

### LDWeaver detects novel interactions between sites associated with virulence in *E. faecalis*

*E. faecalis* represents a classical generalist microorganism, with little phylogenetic divergence and limited host specialisation over the population [64]. Few hospital-associated *E. faecalis* clusters have been identified, with some overrepresentation of virulence factors and AMR genes [27,65]. However, these lineages as well as the traits potentially underlying their success predate the modern hospital settings [27,66].

The *E. faecalis* dataset analysed using LDWeaver comprised 2027 isolates (accessions available in supplementary information) aligned against the V583 reference genome [67] using Snippy. After filtering out sites with MAF < 0.01 and gap frequency > 0.15, 85,982 SNPs remained (see supplementary Fig. 6 for the LDWeaver plot panel).

Inspecting the short-range links within the *E. faecalis* population revealed multiple top-ranking links in known enterococcal virulence genes. Particularly, the *elr* operon was represented in several links (see supplementary Fig. 7). The genes in the *elr* operon code for putative surface proteins and their overexpression have been associated with increased virulence and ability to evade host immune defence by resistance to phagocytosis [68]. Three *elr* genes, *elrB, elrC* and *elrR*, were linked to each other, and the positive regulator *elrE* [69] was also linked to an ATP-binding cassette transporter.

Another cluster of links involved *ace*, a widespread virulence gene in *E. faecalis*, coding for adherence factor [70,71]. In addition to a gene coding for a hypothetical protein, it was linked to a bacteriocin-encoding *entV*. This bacteriocin has been shown to act against fungal *Candida albicans* co-infection by the inhibition of biofilm formation [72,73]. Intriguingly, the absence of ace has also been shown to result in reduced biofilm formation *in vivo*, but despite both *ace* and *entV* being partly involved in biofilm-associated enterococcal infections [74], we are not aware of any report of a direct link between the functions of the two. These findings demonstrate the ability of LDWeaver to highlight both known and putative functional links between enterococcal genes. Figure 8 further illustrates that the minor allele haplotypes at the candidate sites under co-selection are not enriched in hospital-associated multi-drug resistant lineages. Given that the ages of these lineages have been estimated as 50-150 years [27], and that they all share the major alleles at these variable sites, the co-selective pressure may have acted more recently in some other ecological setting outside hospitals.

### LDWeaver detects co-selection signals in simulated genomes

The ability to detect short-range links under selection using LDWeaver was assessed under several simulated evolutionary scenarios. All simulations were conducted using SLiM version 4 [75]. Ten subpopulations of bacteria, each comprising 1000 individuals were simulated. Horizontal gene transfer was simulated following the approach in [76] and the mean tract length was set at 500 bp. Mutation and recombination rates were set at 2e-7 and 8e-7, respectively. Migration was allowed between all 10 subpopulations at a rate of 5e-2. The first 200 kb of the ATCC 700669 reference genome was chosen as the ancestral sequence and the 22 longest CDS (> 1500 bp) were chosen as *potential* regions for co-selection and epistasis. The number of chosen CDS were varied across the simulation scenarios **s1** to **s5**.

Three types of mutations were introduced: the majority of bases outside chosen CDS had the mutation type **m1**, which had a slightly deleterious selection coefficient of -1e-5 (nearly neutral). Each chosen CDS was randomly allocated a mutation type **m2** or **m3**, each with a selection coefficient of +1e-2. Carrying a mutation pair of type **m2** and **m3** had an epistatic effect which resulted in reducing the fitness of the individual by 5%. The simulation was continued for 5000 generations and at the end of the run, 1000 individuals from the total population (10% of the population) were randomly sampled for LDWeaver analysis.

In simulation scenario **s1**, all 22 CDS were chosen as targets, which covered 23.8% of the total genome length. Next, the number of target CDS was reduced four times; in each step the four shortest CDS were converted to near neutral (**m1)** mutations. This resulted in 4 scenarios with a descending number of chosen CDS/genome length coverage - **s2**: 18 CDSs/20.8%, **s3**: 14 CDSs/17.5%, **s4**: 10 CDSs/13.7% and **s5**: 6 CDSs/9.5% (see Fig. 5a). Each simulation scenario (**s1**-**s5**) was replicated 100 times, resulting in a total of 500 simulations.

**Figure 5.**
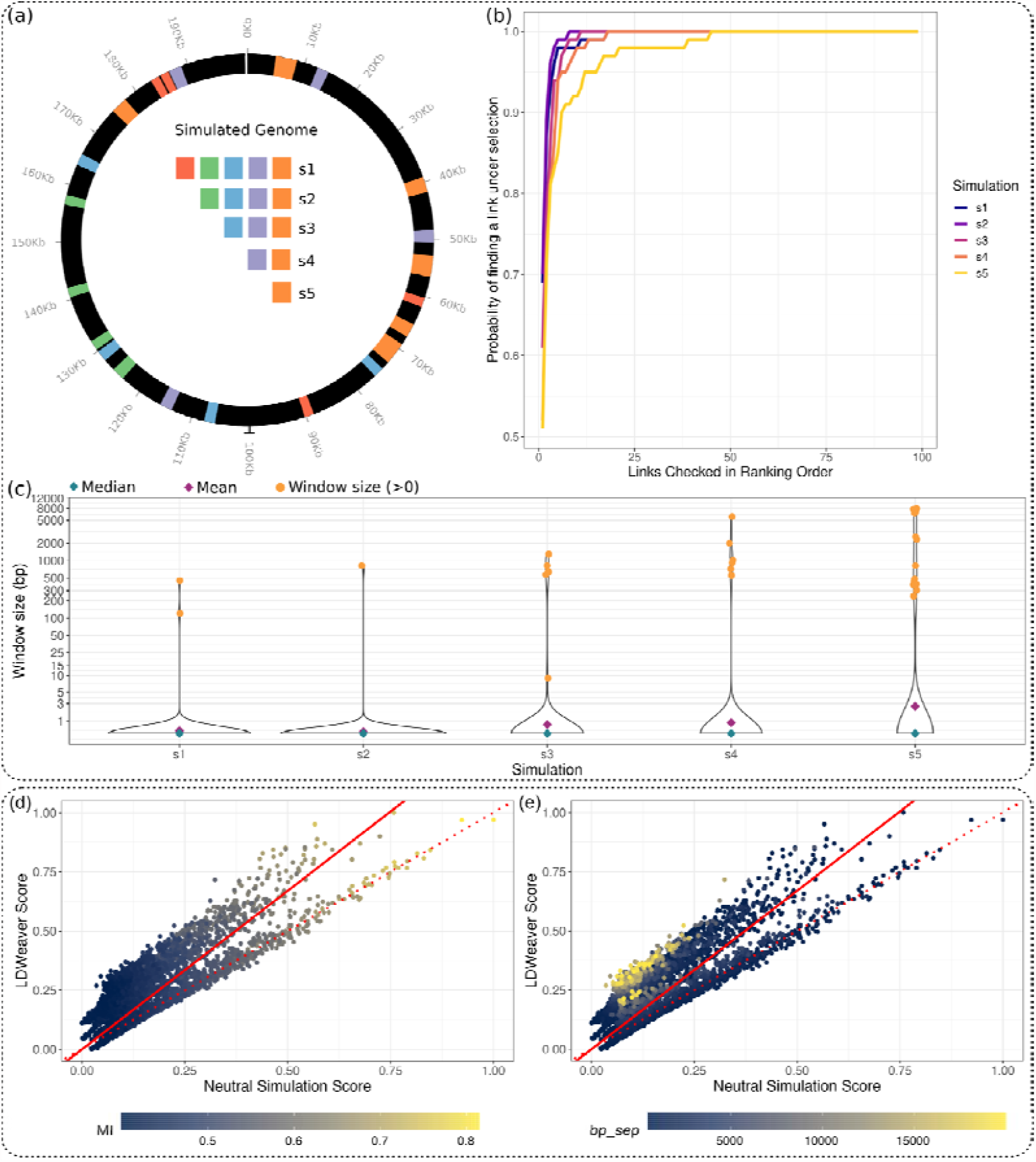
Validation of LDWeaver using bacterial population simulations. (a) Simulated genomic region of 200kb. In each simulation scenario, colour-coded target loci were simulated under selection (e.g., **s1**- all colour coded regions, …, **s5** - orange regions only) - see online methods for details. (b) For each simulation (see legend **s1-s5**), each curve depicts the variation between the number of links checked in ranking order (*x*-axis) and the probability of detecting a target link (i.e., a link from within or between two target regions) (*y*-axis). This probability = 1 when all replicates contain a target link. 97% of replicates from the most challenging s5 scenario contain a target link within the top 20 ranked links. It is noted that in **s5**, <10% of the genome is under selection and only the two orange regions proximal to 70 kb in (a) can contribute to between-region short-range target links. (c) The *y*-axis shows the distance between the top 5 ranked links and a target region for each simulation scenario (*x*-axis). The vast majority of replicates in all scenarios contain a target link in at least one of the top 5 (blue diamond - median = 0, purple diamond - mean < 3 and increases from **s1** to **s5**). Each orange dot corresponds to a replicate that does not contain a target link and its *y*-axis value shows the window size required to get the link to fall within a target region (i.e., the distance from the further site in the link to its closest target region). For example, two replicates in s1 do not contain a target region link in the top 5 and the closest site in each link is approx. within a 100 and 500 bp window, respectively. (d, e) The *y-axis* shows the LDWeaver score, which is the ranking scaled as a continuous variable between 0 and 1, versus the neutral simulation score (*x*-axis) for short-range links from the *Campylobacter jejuni* dataset. The red lines show x=y (dotted) and the best fit (solid). Colour shading indicates (d) MI values and (e) bp-sep. Links with high MI values (>0.5) are ranked similarly using both approaches (d), but links that are far apart are given higher scores by LDWeaver compared to neutral simulations (e).

Within the top 20 ranked short-range links, at least one ‘target link’ (i.e., a link from within or between two chosen CDS regions) appeared in all and 97% of replicates in scenarios **s1**-**s4** and the most challenging **s5**, respectively (See Fig. 5b). Additionally, analysing the top 5 links revealed that for scenarios **s1-s5**, on average 70.4%, 62.6%, 61.8%, 52.4% and 49.8% were target links and in the vast majority of replicates, a target link was included in the top 5 (See Fig. 5c).

Finally, there was strong agreement in link ranking between using the LDWeaver model-free algorithm and the computationally expensive neutral model-based approach for the *C. jejuni* dataset (see Figs. 5d and 5e). This was particularly true for high-LD links (MI > 0.5), which indicates that the process of ranking high-LD links is robust to the choice of method.

## Discussion

Bacterial population genomics research is rapidly moving towards an era where hundreds of thousands of whole-genome sequences will be available for many species. These data represent an unprecedented opportunity to seek signals of selection in natural populations and to unravel genomic clues for adaptation under changing ecology, which can contribute towards improved understanding about evolution, dissemination and maintenance of antibiotic resistance and virulence traits. Existing GWES methodology has already enabled discoveries of co-selection affecting polymorphisms across distant genomic sites in a variety of human pathogens [13–15]. By extending GWES to joint screening of polymorphisms within close proximity to each other, we increase the potential of data-driven molecular discovery for bacterial populations, which are particularly challenging for GWAS due to the difficulty of large-scale measurement of traits.

Applying our methodology across a diverse spectrum of bacterial species: *S. pneumoniae, C. jejuni, E. coli* and *E. faecalis*, has revealed fresh insights into genomic interactions between variants within close proximity to each other. Notably, our results were obtained without a reliance on accurate phenotype data and were otherwise undetectable via conventional genome-wide association (GWAS) methods as they would typically be discarded due to the influence of short-range linkage disequilibrium. Our results included findings associated with host range, antibiotic resistance, virulence and immune evasion. Furthermore, our top results often represented links from genomic islands: *tfoX* in *S. pneumonaie, cdtABC* in *C. jejuni*, capsular locus in *E. coli* and *elr* genes in *E. faecalis*, all of which are self-contained units that may evolve differently to the rest of the chromosome due to reduced functional integration. While the identified novel interactions between these loci require experimental validation, our approach drastically reduces the number of pairwise relationships that need to be considered. Recent advances in the computational protein structure prediction could further reduce the necessity for time-consuming wet-lab experiments by rapidly considering the impacts of the identified intragenic interactions on the resulting protein structure.

A potential target for further development of genome-wide short-range LD analysis is to consider genomic variation beyond reference-based core genome alignments. With the increasing availability of long-read based assemblies of chromosomes and plasmids, it would be attractive to consider detection of co-selection both within plasmids and between plasmid and host chromosome polymorphisms, to potentially uncover either compensatory evolution or pre-adaptation to stable carriage of particular plasmids [77]. Given its existing and future potential for enabling molecular discoveries, we anticipate that GWES will continue to attract a wide interest from both methodological and applied perspectives.

## Online Methods

### Measuring LD

By default, LDWeaver removes sites with MAF < 0.01 and gap frequency > 0.15. Sites with ‘gap’ as the second most common allele are also discarded from the analysis by default, but LDWeaver has a filtering option called “relaxed” that retains these sites in the analysis. This option could be particularly useful for alignments with a limited number of SNPs, as gaps can reflect insertions or deletions with functional effects [78].

Following SpydrPick [12], we use MI to measure LD. The pairwise MI between two sites (modelled as discrete random variables) is given by:

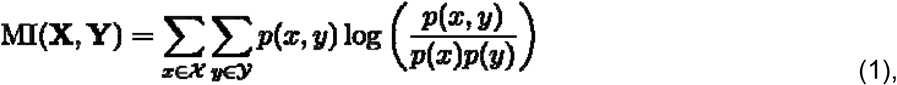

where **X** and **Y** denote two sites with alleles ***x* ∈ 𝒳** and ***y* ∈ 𝒴**, respectively. For SNP data, the alphabet comprises nucleotides A, C, G, T and the gap character N. Here, ***p(x***,***y)*** denotes the joint probability of **X = *x*** and **Y = *y*** and ***p(x), p(y)*** are the corresponding marginal probabilities which are estimated from count data. Let denote the number of sequences in the alignment, then:

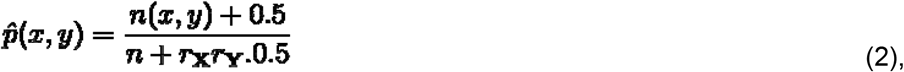

where ***n*(*x***,***y*) =** | **S**_***xy***_| is the number of sequences with **X = *x*** and **Y = *y*, r**_**X**_ **=** | **X** | and **r**_**Y**_ **=** | **Y** |.

Strong population structure in bacteria presents a problem for analysis [78]. We adopt a widely-used sequence-reweighting approach [10–12,79,80]. The weight ***w***_***i***_ **∈ [1/n**,**1]** for sequence ***i*** is computed as the reciprocal of the number of sequences with mean per-site Hamming distance **< *t***, where ***t*** is a dataset-dependent threshold that typically satisfies ***t* ∈ (0.1**,**0.25)**. This population structure correction is applied to the LD structure estimation by substituting effective counts in eq (2) given by 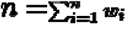 and 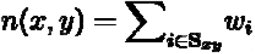.

In practice, the MI computation process is optimised using a sparse matrix representation and performed in blocks of SNPs [81], which can be feasible even in systems with relatively low memory. LDWeaver requires genome-wide short-range links to be retained in memory (see below), but most long-range links with low MI values are discarded after processing each SNP block. The user specifies an approximate number of long-range links to be retained for downstream analysis.

### Modelling short-range links

By default, SpydrPick [12] uses **𝒮 = 10** kb as the threshold for defining a short-range link, but in LDWeaver we chose **𝒮 = 20** kb as the default threshold to better capture the region of rapid LD decay. The user can adjust this parameter as required.

Additionally, LDWeaver accounts for genome-wide variation in local LD patterns, which can arise due to varying mutation and/or recombination rates, by introducing a clustering and segregation step. First, for each coding sequence (CDS) segment in the annotations file, the per-site number of mismatches between the CDS and its reference sequence (i.e. the persite Hamming distance) is computed. Next, the CDS are clustered using the k-means algorithm [82]. Since CDS regions exclude intergenic regions, each intergenic SNP is manually merged into the cluster closest to its genomic location.

For comparison purposes we model the rapid LD decay of short-range links using computationally demanding neutral simulations. For each cluster, we used mcorr [83] to estimate the parameter triplet: mutation rate, recombination rate and the recombination tract length. For each parameter triplet, 100 neutral replicates were generated using bacmeta [84]. In each simulation 20 subpopulations of bacteria, each comprising 1000 individuals with a 200 kb genome were included to generate a sufficiently large and diverse alignment. Each simulation was performed for 20000 generations to ensure convergence and then 1000 individuals were sampled. The mean LD decay was directly estimated from this neutral data as a function of bp-sep.

LDWeaver directly models the LD-decay using the genomic alignment, separately for each cluster. At each discrete bp-sep, the 95th percentile (**q95**) MI value is extracted from the genomic data. Next, the linear model: **log (q95) ∼ log (bp_sep)** is fitted and the exponent of the fitted values of this model (i.e.,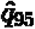) is chosen as the bp-sep dependent short-range background-LD threshold. The 95th percentile is chosen to maintain a conservative threshold such that a sufficiently large number of links are retained for downstream analysis.

### Outlier calling and link ranking

Outlier calling is performed at each discrete bp-sep value. Let ***d* ∈ [1, 𝒮]** denote the bp-sep of interest, let **L(*d*)** denote all the links that are ***d*** bp-sep apart and let **L ∗ (*d*)** denote the subset with 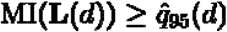. First, the model: 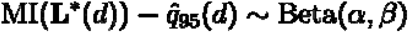 is fitted [85]. Afterwards, a short-range *p*-value is computed for all **L ∗ (*d*)**. Once the outlier calling is performed **∀*d* ∈ [1, 𝒮]**, this **−*log***_***10***_ **(p)** value (referred to as ‘srp’ in LDWeaver to denote shortrange p-value) is used to assign a rank for each link. All links in **L(*d*)** that do not belong to **L ∗ (*d*)** (i.e., links with 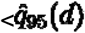) are discarded from downstream analysis and the user can choose a srp cut-off value to discard links (default cut-off ***p* = 0.001**).

Since 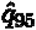 values vary over genome clusters, links from separate clusters are ranked based on the ‘local’ background-LD decay. When a link spans two genomic clusters, the maximum of the two srp values is used for ranking and downstream analyses.

### Filtering indirect links

Because of its success in inferring gene expression networks [86] and its utility in SpydrPick, we added ARACNE as a filtering step to the LDWeaver pipeline to overcome the inability of pairwise methods to distinguish “direct” associations. For a dataset with 100 000 SNPs, MI values will be computed for approximately 5 billion unique links. When performing GWES, the interest is typically focussed on links with high MI values (i.e., with larger than normal linkage). However, many of these links will be driven by the same underlying association. Identifying the causal link is extremely challenging, especially for bacterial data due to factors such as clonality and genome plasticity [78].

Given multiple links that can be explained by the same underlying causal effect, we use ARACNE to retain only the link with the strongest signal (called a “direct” link). This can reduce the number of links retained by several orders of magnitude, greatly reducing the manual curation task. ARACNE scans the entire LD landscape and a link (**X**,**Y**) is considered to be indirect if **MI(X**,**Y) ≤ min**[**MI(X**,**Z), MI(Z**,**Y)]** for any SNP Z. A detailed explanation of the ARACNE algorithm and this specific filtering step is available in [12].

### Detecting outliers in long-range links

Long-range (bp-sep > **𝒮**) background LD varies little with bp-sep, so the genome clustering step that was used to model short-range links is not required. We follow an approach similar to that of SpydrPick to determine a background-LD threshold [12].

After long-range MI, LDWeaver computes the outlier threshold defined by Tukey [87]: **𝒯 = Q**_**1**_ **+ 1.5 × IQR**, where **IQR** = **Q**_**3**_ **− Q**_**1**_ and **Q**_**1**_ and **Q**_**3**_ are the first and third quartiles, respectively. Links with MI **> 𝒯**_**1**_ are directly ranked based on the MI value. In cases where < 5000 links surpass **𝒯**_**1**_, LDWeaver will pick the top 5000 links based on MI value for downstream analysis. Similar to the short-range analysis, the top 5000 links are chosen as a conservative threshold such that a wide range of links are available for downstream analysis (see below).

While the Tukey outlier detection approach is robust against extreme values, it appears simplistic compared to a permutation approach, which is typically preferred to determine a threshold for these types of analyses. However, permutation testing is undesirable here for at least two reasons. Firstly, this threshold has limited impact on the long-range GWES analysis itself. While an outlier is defined based on whether its MI value passes this particular threshold, it has no bearing on link ranking itself. Downstream analysis and the eventual manual curation can be performed on the highest-ranked subset of ARACNE direct links without requiring a threshold. Secondly, due to the high degree of LD observed in bacteria, the background MI values computed from the observed MSA could be larger than the null MI values computed from a label-shuffled model. Therefore, the computed permutation threshold could potentially be too conservative.

While LDWeaver can perform GWES analysis on both short- and long-range links, it is also fully compatible with the SpydrPick output for long-range link analysis. For a dataset that has already been analysed using SpydrPick, users have the option to first perform only the shortrange analysis in LDWeaver, then present the SpydrPick output as input to LDWeaver to perform the downstream analyses of long-range links.

### Downstream analyses and visualising putative links and sites

We have integrated several powerful R visualisation tools into LDWeaver [88], [89]. In order to prioritise and perform wet-lab based modelling and validation, it is helpful to understand the functional annotations of the SNPs involved in high-MI links. Therefore, functional annotations are added to all such SNPs using SnpEff [28]. Additionally, LDWeaver classifies each SNP as non-synonymous, synonymous or intergenic. The non-synonymous versus synonymous distinction is based on the most common allele at multi-allelic sites. Based on this classification, LDWeaver generates an additional output comprising, by default, a set of 250 *top ranked links* by discarding links between two synonymous sites.

GWES Manhattan plots were introduced to visualise the distribution of MI as a function of bp-sep [12]. LDWeaver generates the long-range GWES plot introduced in SpydrPick, along with two plots for short-range links. The first short-range plot is similar to the long-range GWES plot but with points shaded according to the srp value. The second plot shows the segregation of links into genomic clusters. While these plots are less informative compared to Manhattan plots in genome-wide association studies (GWAS) due to the lack of genomic positions, they can be useful to assess the LD-decay fit between 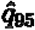 and the genomic data.

LDWeaver also generates a linear tanglegram using the R package ChromoMap [90] to indicate the genomic positions of *top ranked links* in the short range. These tanglegrams span the whole genome and are broken down into segments for improved utility. Additionally, LDWeaver generates a figure depicting a genome-wide overview of the LD structure in the dataset. First, a sparse, SNP-level LD matrix is created using the MI values of the saved link data. Then, an averaging kernel is applied to reduce the dimensions of the matrix to approximately **1000 × 1000** [91]. The resulting matrix is plotted in the form of a heatmap with SNP positions as *x* and *y* labels [92]. This bird’s-eye view can be useful in some analyses to quickly identify high-LD regions and blocks.

Additionally, LDWeaver generates a network plot for *top ranked links* using the R package ggraph [93]. The gene-level regions of each site are extracted from the annotations file and the links between these regions are coerced into a network [94]. To reduce clutter in the network plot, edges with only 1 link between nodes are removed. Afterwards, any nodes with no edges between them are also dropped from the plot. The edges are coloured to depict the number of links between genes. Furthermore, the edge width and transparency are also moderated to reflect the MI value of the highest ranked link between the two regions. Furthermore, LDWeaver provides the option to generate a similar gene network for any chosen gene. The user can decide whether to use short-range, long-range or both link lists to generate this plot.

Using a user-provided phylogeny, LDWeaver can generate a tree-plot of user-determined putative sites and phenotypes. This type of plot is inspired by some of the visualisation options available in Microreact [95] and has also been widely used in the previous GWES literature [11,15]. The phylogeny will be midpoint rooted by default [96,97] and the SNP data, user provided phenotypes are sorted in the same order as the phylogeny. Finally, the plot is generated using the R package ggtree [98].

Finally, LDWeaver generates the output required to dynamically visualise links using the R package GWESExplorer (https://github.com/jurikuronen/GWES-Explorer). This Node.js based shiny app can be used to generate the GWES Manhattan plot, circular tanglegram and the tree-plot for an arbitrarily chosen subset of putative links.

## Supporting information

Data Accessions

## Software Availability

LDWeaver v.0.1.0 is available as an R package under a GNU General Public License (Version 3) on GitHub (https://github.com/Sudaraka88/LDWeaver) and the source code is available on Zenodo (https://zenodo.org/record/8201753).

## Runtime

A complete LDWeaver analysis with default parameters on a dataset comprising 2000 sequences with 80,000 SNPs on average requires 4 756 seconds (∼80 minutes) on a computer with 32GB of ram and 10 parallel CPU cores running R version 4.2.2 with openBLAS v0.3.21 support.

## Funding Information

J.C. and GTH were funded by NFR grant no. 299941. S.M., N.M., S.A.-A., R.A.G. and A.K.P. were funded by the AMR grant from Trond Mohn Foundation and S.M., N.M. and S.A.-A. were additionally funded by Marie Skłodowska-Curie Actions (Grant 801133). NJC was supported by a Sir Henry Dale Fellowship, jointly funded by Wellcome and the Royal Society (grant no. 104169/Z/14/A; https://wellcome.org/; https://royalsociety.org/).

## Supplementary Information

Dataset accessions available at: <u>datasets_LDW.csv</u>

## Supplementary Figures

**Supplementary Figure 1.**
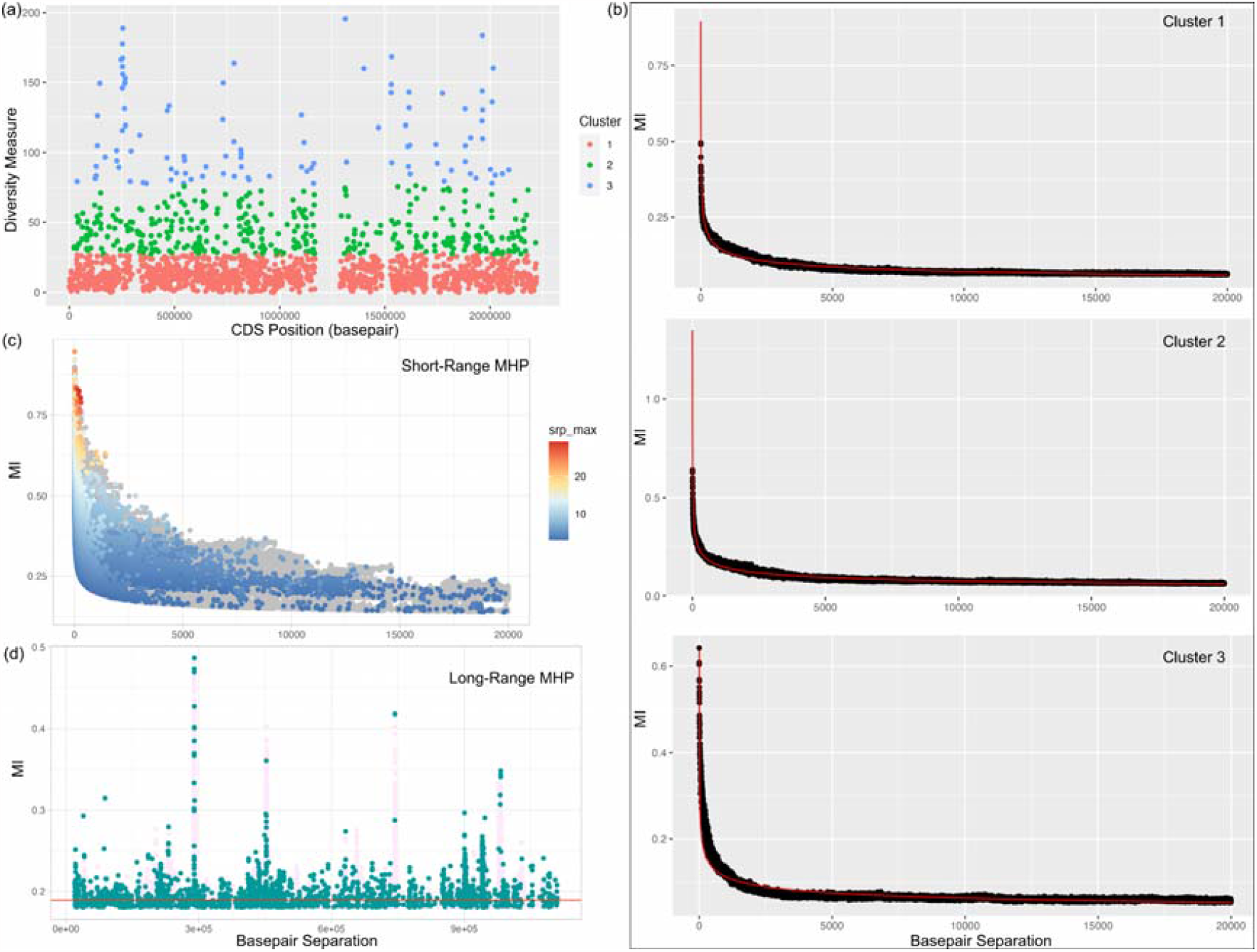
Panel of LDWeaver plot for the *Streptococcus pneumoniae* Maela dataset aligned using the ATCC700669 reference genome. (a) Diversity measure of each CDS in the reference genome annotation. K-means clustering was performed to allocate each CDS to one of three clusters (colours) based on the diversity. (b) For each cluster, the panel shows the LD-decay of the empirical 95th percentile MI value at each bp-sep (black dots) and the fitted values 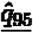 (red line). (c) In the short-range Manhattan plot, the *x*-axis shows the bp-sep and the *y*-axis shows the MI value. Each coloured point corresponds to an outlier link in the dataset. Links are coloured according to the srp value computed using the background-LD distributions in (b). When a link stems from two clusters, srp is computed for both backgrounds and the maximum value is used. Points in ash colour are ARACNE indirect links. (d) The long-range Manhattan plot conveys similar information to (c) and the red horizontal line shows the Tukey outlier cutoff.

**Supplementary Figure 2.**
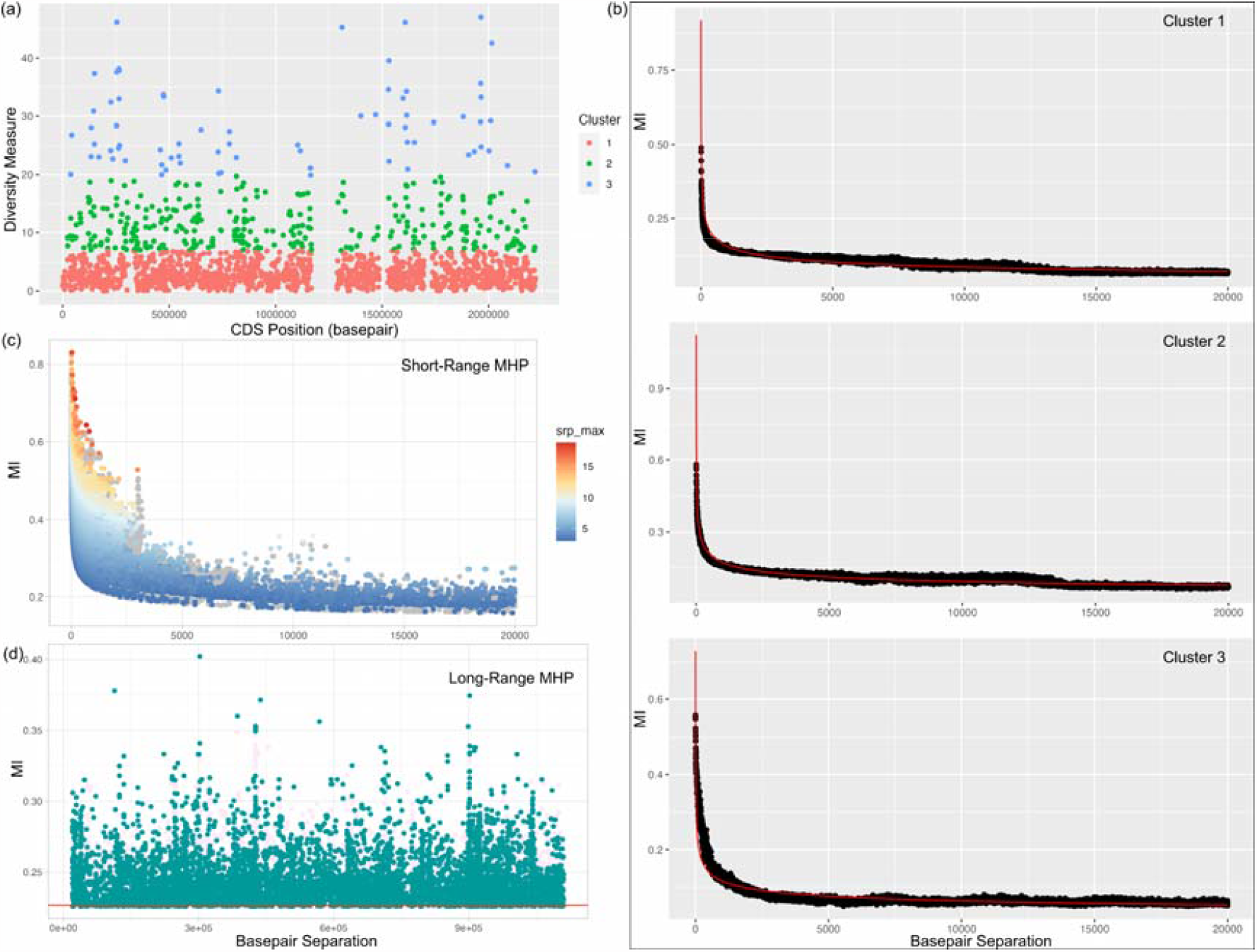
Panel of LDWeaver plot for the *Streptococcus pneumoniae* Massachusetts dataset aligned using the ATCC700669 reference genome. See Supplementary Figure 1 caption for figure details.

**Supplementary Figure 3.**
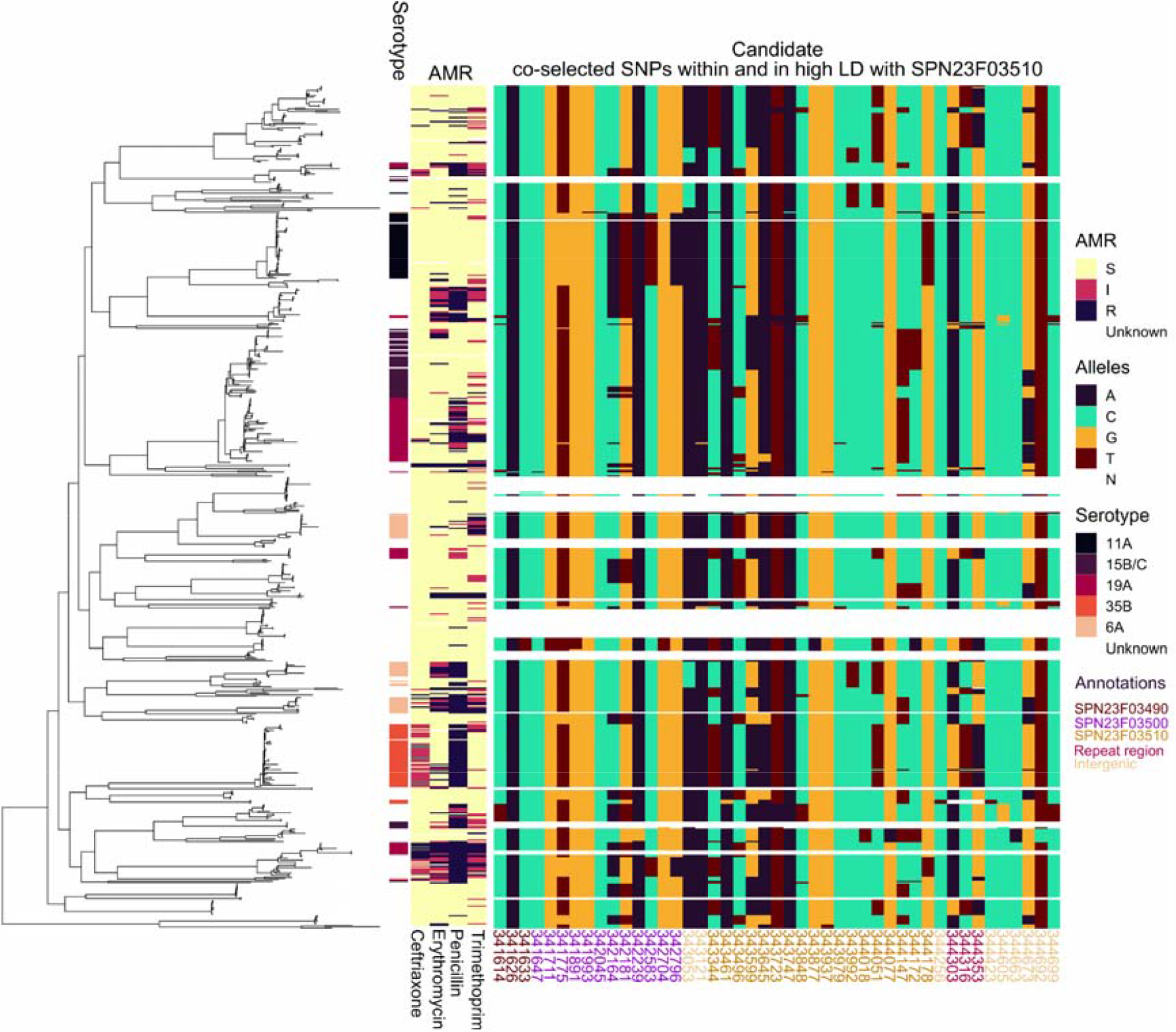
Overview of the allelic variation within SPN23F03490 - SPN23F03510 region in the Massachusetts *S. pneumoniae* dataset. (a) The phylogenetic tree (n = 616) was estimated using FastTree 2 [99] and the 5 bars immediately to the right denote the serotype, and antimicrobial resistance data for ceftriaxone, erythromycin, benzyl penicillin and trimethoprim, respectively (keys are shown on the right and the AMR colour shadings indicate S - sensitive, I - intermediate and R - resistant). Allelic variation is shown by the heatmap to the right and the corresponding key shows the colours for nucleotides with N indicating gap. SNP positions below the heatmap are based on the FM211187 reference and are colour coded to indicate the annotation (key on the right).

**Supplementary Figure 4.**
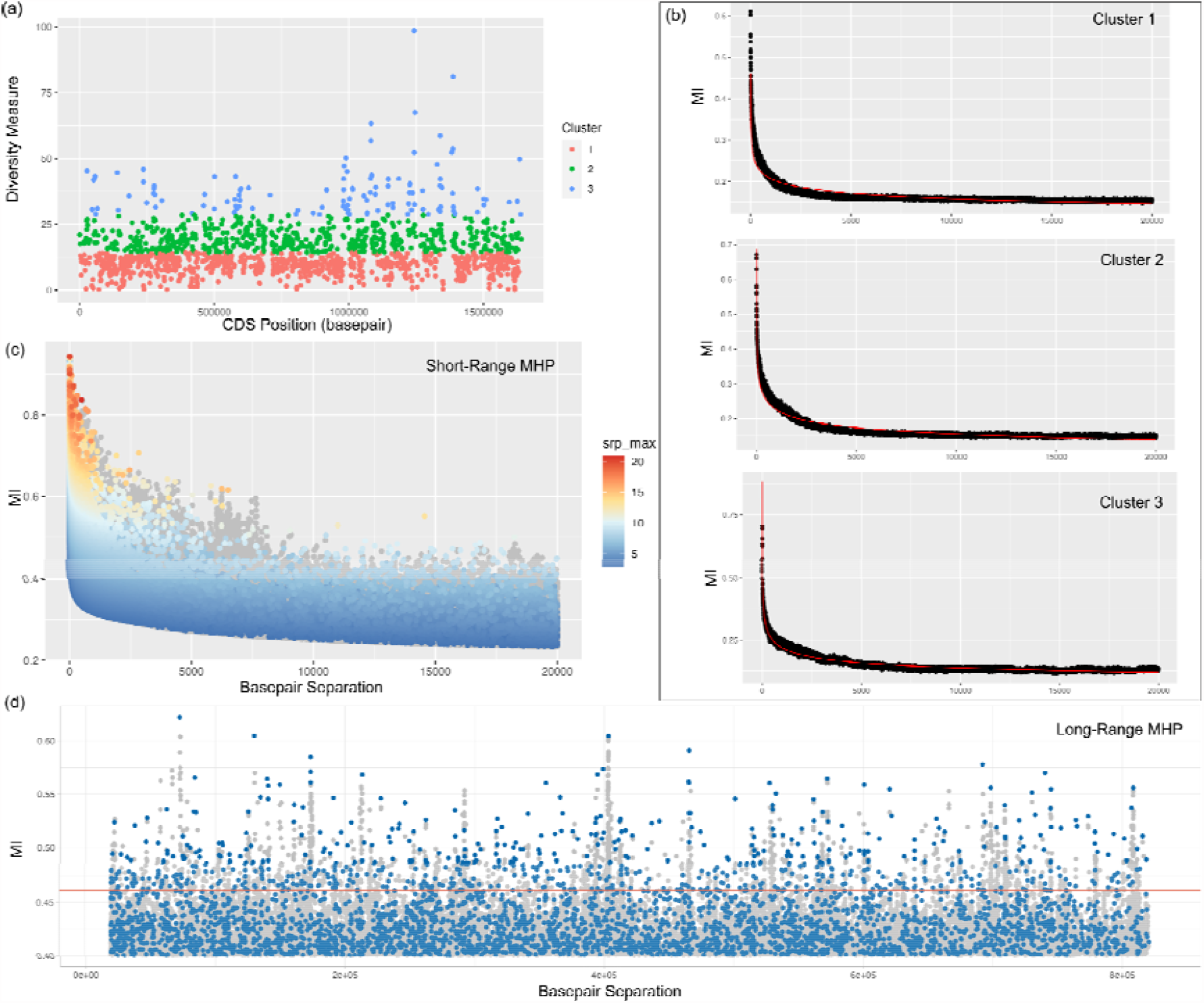
Panel of LDWeaver plot for the *Campylobacter jejuni* dataset aligned using the NCTC 11168 reference genome. See Supplementary Figure 1 caption for figure details.

**Supplementary Figure 5.**
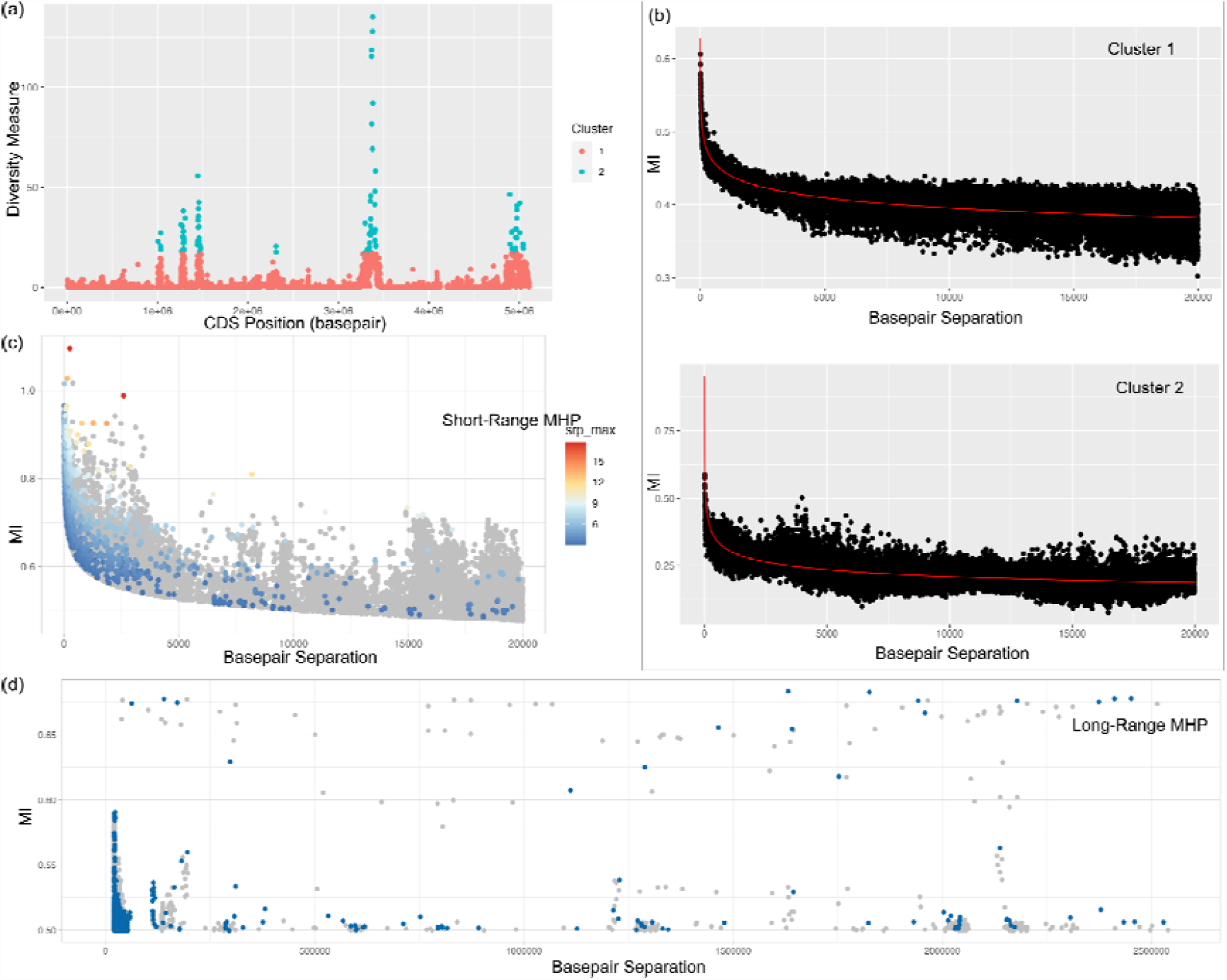
Panel of LDWeaver plot for the *Escherichia coli* dataset aligned using the EC 958 reference genome. See Supplementary Figure 1 caption for figure details.

**Supplementary Figure 6.**
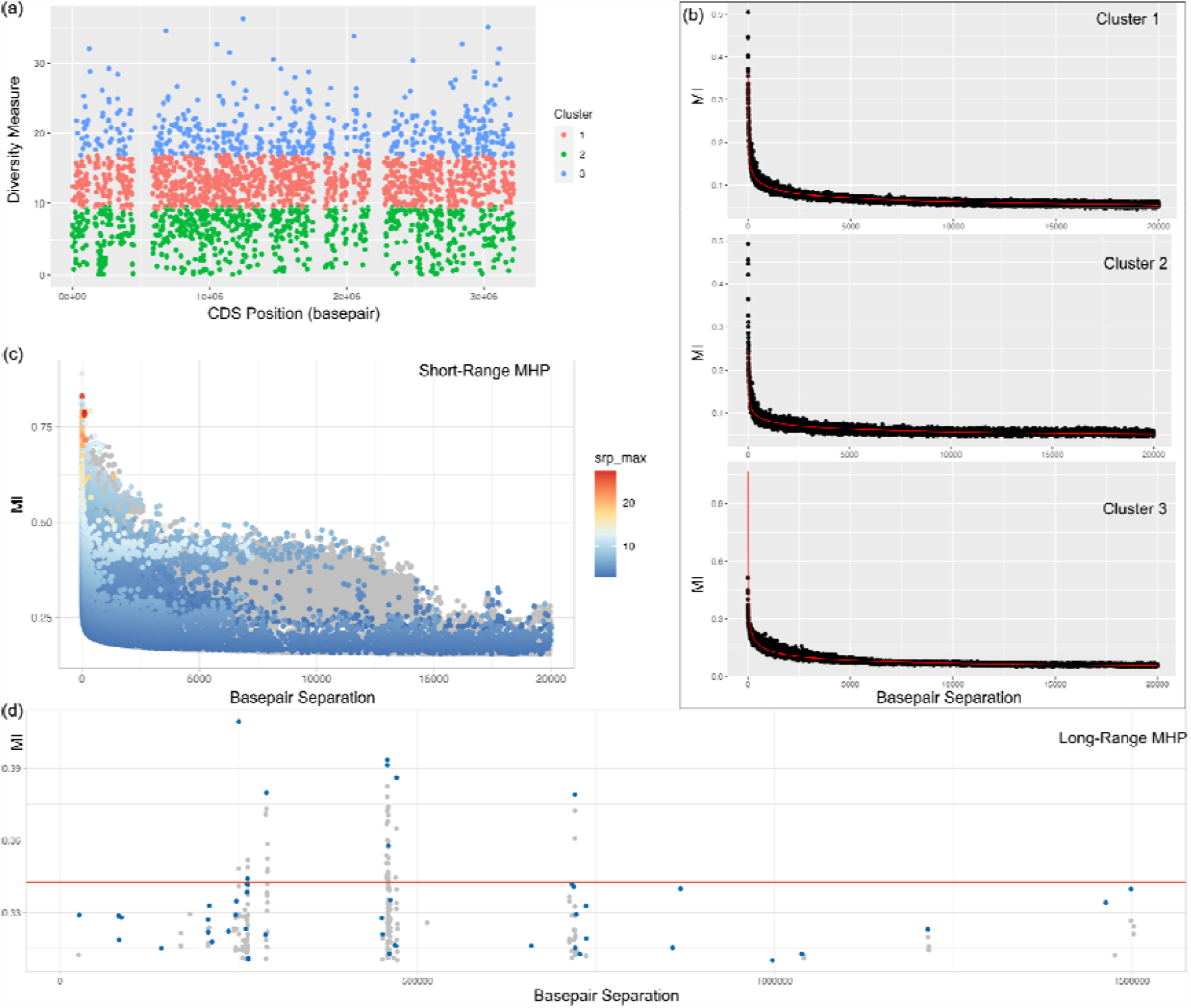
Panel of LDWeaver plot for the *Enterococcus faecalis* dataset aligned using the V583 reference genome. See Supplementary Figure 1 caption for figure details.

**Supplementary Figure 7.**
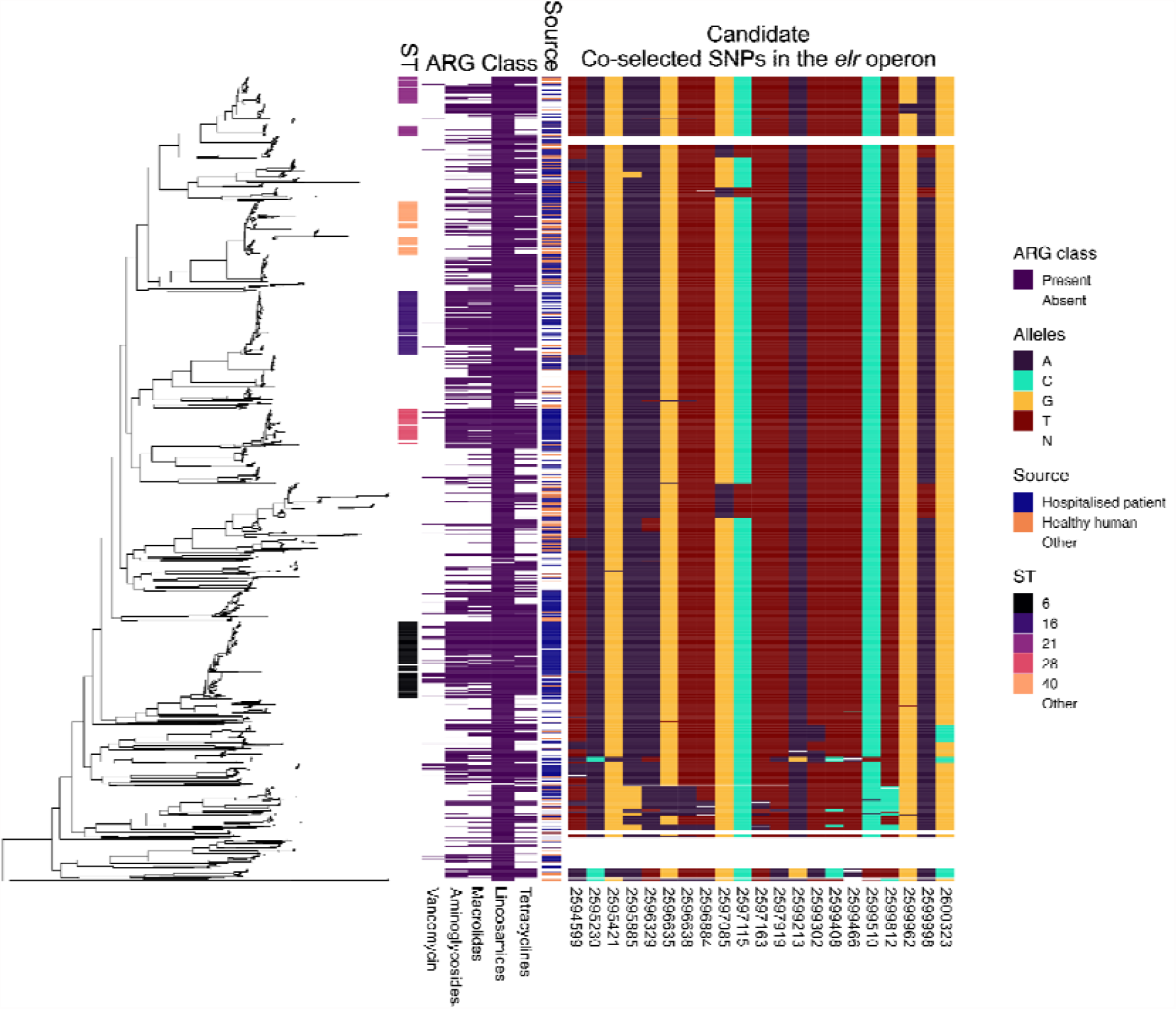
Approximate maximum likelihood phylogeny for the coregenome alignment, estimated using FastTree 2 [99], aligned to panels of metadata and a subset of alleles in the *elr* operon. From left to right, panels show, from left to right ST - sequence type, ARG class - Antimicrobial Resistance Genes (see below for each ARG), Source of isolate and the Allele distribution for 21 SNPs (see below for genomic position as per the V583 NC_004668.1 genome). All SNPs shown in the panel were detected as short-range top ranked links by LDWeaver.

## Notes

https://github.com/Sudaraka88/LDWeaver

https://zenodo.org/record/8201753

https://unimelbcloud-my.sharepoint.com/:x:/g/personal/sudaraka_mallawaarachchi_unimelb_edu_au/EdF0R3HClHJApsvu7SpoyFMB7urh-NAPTgGkiuftP9cWNA?e=5tjg6w

https://cloudstor.aarnet.edu.au/plus/s/KBRnIt1H6XZ2XFR

### Competing Interest Statement

The authors have declared no competing interest.

### Summary of Updates

Added author ORCID and github link to abstract

